# Electrostatic Collapse of Intrinsically Disordered Acid-Rich Protein Is Sensitive to Counterion Valency

**DOI:** 10.1101/2025.07.09.663830

**Authors:** Barbara P. Klepka, Radost Waszkiewicz, Michał Wojciechowski, Agnieszka Michaś, Anna Niedzwiecka

## Abstract

Intrinsically disordered proteins (IDPs) respond sensitively to their ionic environment, yet the mechanisms driving ion-induced conformational changes remain incompletely understood. Here, we investigate how counterion valency modulates the dimensions of an extremely charged model IDP, the aspartic and glutamic acid-rich protein AGARP. Fluorescence correlation spectroscopy and size exclusion chromatography reveal a pronounced, valency-dependent reduction in its hydrodynamic radius, with divalent cations (Ca^2+^, Mg^2+^) inducing collapse at much lower activities than monovalent cations (Na^+^, K^+^). Molecular dynamics simulations, direct sampling, and polyampholyte theory quantitatively capture the Debye–Hückel screening by monovalent ions but not the enhanced compaction driven by divalent ion binding. Circular dichroism spectroscopy shows that compaction occurs without secondary structure formation. Our results demonstrate a structure-free electrostatic collapse and suggest that specific chelation of divalent ions by disordered polyanionic protein chains is a key mechanism regulating IDP compaction, with implications for understanding their behavior in biologically relevant ionic environments.

**TOC GRAPHIC:** 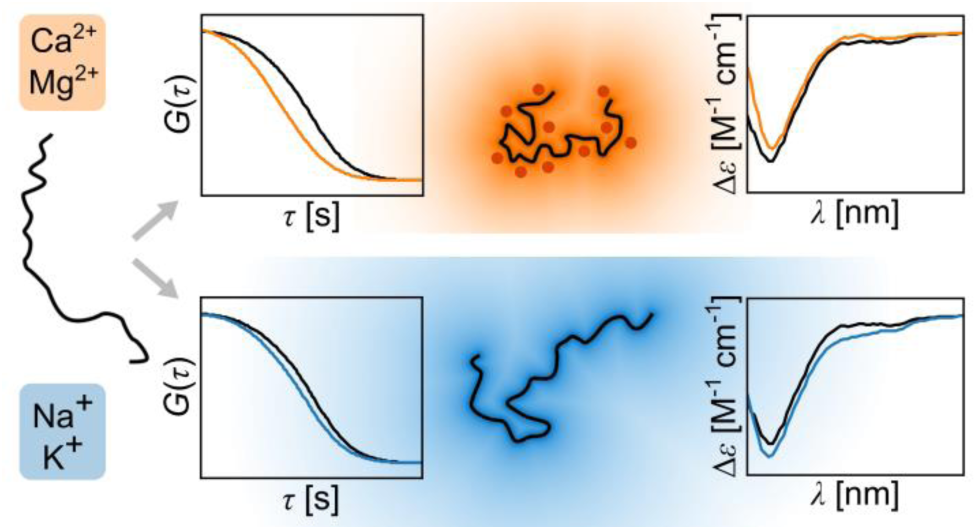

---

Intrinsically disordered proteins are essential regulators in diverse biological processes, including cellular signaling,^1,2^ gene expression,^3^ or biomineralization.^4^ Despite their high abundance,^5^ our understanding of the determinants of their conformational ensemble equilibria and function remains incomplete.^6^ Unlike natively folded proteins, which adopt a singular three-dimensional structure, IDPs exhibit remarkable conformational dynamics and are best studied as rapidly fluctuating ensembles of conformations. IDPs are typically composed of low-complexity sequences, enriched in polar and charged residues and depleted in hydrophobic ones.^7^ Thus, their behavior is often described within the frameworks of polyampholyte and polyelectrolyte theories.^8–15^ The apparent dimensions of IDP conformational ensembles are usually quantified using the radius of gyration (*R_g_*) or hydrodynamic radius (*R_h_*). These parameters have been shown to vary not only with the protein sequence,^16^ but also in response to environmental conditions,^17^ such as temperature,^18^ pH,^19^ osmolality^20^ or ionic strength.^21–27^ In particular, the effect of solution ionic composition has been examined; in the case of monovalent salts, classical polymer models have demonstrated a high level of agreement with experimental observations of changes in the dimensions of charged proteins due to electrostatic screening.^21,23^

Although the function of certain IDPs depends on their interactions with divalent ions, particularly the acid-rich proteins involved in biomineralization,^28–31^ the effects of these ions on IDP dimensions remain poorly understood. Most prior research has focused on monovalent salts,^21–23^ oversimplifying more complex relationships. Consequently, protein-salt interactions are reduced to the effect of electrostatic screening, precluding the ability to address specific counterion recognition.

To address this knowledge gap, we examined the effects of mono- and divalent cations on the hydrodynamic dimensions of the polyanionic aspartic and glutamic acid-rich protein (AGARP)^32^ from the coral *Acropora millepora* of the Great Barrier Reef. AGARP is an extremely charged, natively unstructured protein^31^ belonging to the family of coral acid-rich proteins (CARPs), which are found in stony corals.^32,28^ These proteins are secreted at the organism–seawater interface, where they are thought to regulate the formation of CaCO_3_ skeletons.^33^ Ca^2+^ and Mg^2+^ ions also play a central role in regulating this process, since they directly contribute to aragonite deposition in corals.^34^ AGARP was recently shown to undergo liquid–liquid phase separation driven by the interactions with Ca^2+^ under molecular crowding conditions.^31^ The AGARP polypeptide chain is composed of 506 amino acid residues, including a total of 212 negatively charged and 65 positively charged residues (**Figure S1**, **Table S1**). At pH 8.0, AGARP has a net charge of −148 *e* per molecule and is devoid of folded domains.^31^ Thus, AGARP serves as an excellent model protein, with a linear charge density of –0.3 *e*/residue, which is comparable to that of prothymosin α (–0.4 *e*/residue). However, AGARP has a much longer chain (506 *vs*. 111 residues), resulting in greater relative size changes due to the electrostatic self-repulsion.

To elucidate the influence of the environmental electrostatic conditions on the conformational properties of AGARP, we employed a combination of several experimental, numerical, and theoretical approaches. First, we experimentally determined the hydrodynamic radius of AGARP as a function of increasing concentrations of mono-(NaCl, KCl) and divalent (MgCl_2_, CaCl_2_) salts using fluorescence correlation spectroscopy (FCS)^35,36^ and size exclusion chromatography (SEC) (for details see Supplementary Materials). Second, we computed the *R_h_*values using Minimum Dissipation Approximation (MDA)^36,37^ based on conformers generated by coarse-grained molecular dynamics simulations (CG MD)^38,39^ and direct conformational sampling (Globule-Linker Model, GLM).^36^ Third, we compared the experimental and numerical results against the theoretical polymer model of Higgs and Joanny^40^. Fourth, we used circular dichroism (CD) spectroscopy to assess whether the counterion-induced compaction of AGARP could be attributed to changes in its secondary structure. We concluded that, surprisingly, the electrostatic collapse of AGARP occurs without protein folding. Finally, we showed that the decrease in the *R_h_*values conforms to the classical Debye-Hückel theory for monovalent cations. The response of the highly charged protein chain to the presence of divalent salts, by contrast, is additionally enhanced through its specific interactions with the counterions.

The experimental backbone of the presented investigation is formed by measurements obtained using two complementary methods for determining *R_h_*: FCS, which is the only technique that allows for direct measurement of self-diffusion, and SEC. For all experiments, AGARP was purified as previously described^31^ and treated with Chelex^®^ 100 resin to remove divalent ions. The protein labeled with AF488 (Lumiprobe) was used for FCS, and the unlabeled protein was used for SEC.

FCS measurements were performed essentially as described previously,^35,36^ in 30 µL droplets at 25 °C using a titration approach, where aliquots of salt stock solutions in water were sequentially added to a 100 nM protein sample in 10 mM Tris/HCl, 5% glycerol, pH 8.0 (TG buffer), while carefully monitoring evaporation. The evaporation rate was determined prior to titrations by measuring the FCS amplitude for AF488 freely diffusing in the buffer solutions with given salt concentrations. This ensured that the droplet volume was known, which enabled the calculation of the actual salt concentration and the determination of the sample viscosity. A two-component model of 3D diffusion that accounts for residual amounts of free dye in protein samples was fitted to all measurements. The model included a fixed triplet state relaxation time for AF488 covalently attached to the protein of 2.4 μs as the average lifetime, determined from multiple independent experiments. This allowed us to extract diffusion times and determine the exact *R_h_* of the protein under different conditions.^37^ Representative results of the FCS measurements are shown in **Figure 1**.

**Figure 1.**
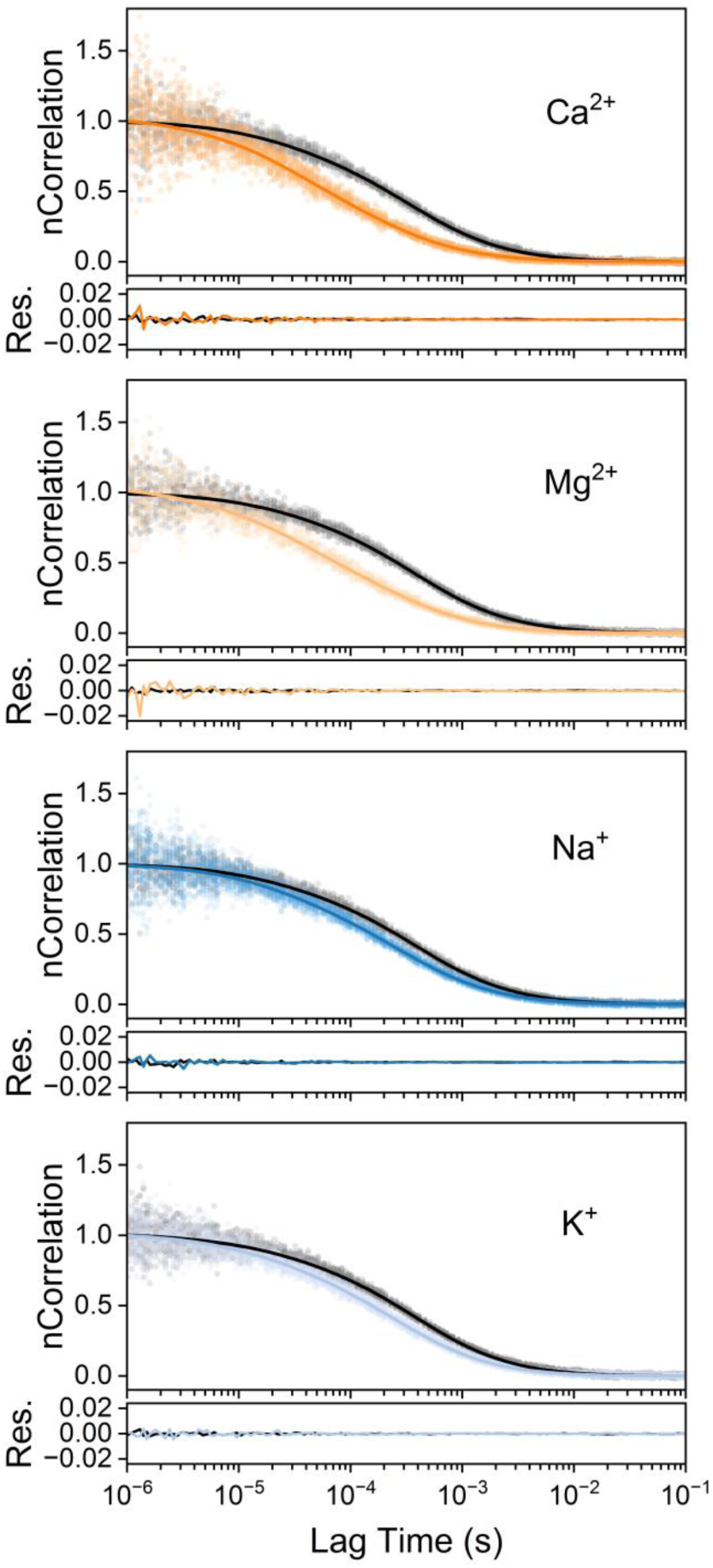
Effects of cations on the diffusion of AGARP, measured by FCS. Examples of normalized FCS autocorrelation curves (solid lines) from global fitting to FCS data (translucent points) along with raw fitting residuals for 150 mM CaCl_2_, 130 mM MgCl_2_, 490 mM NaCl, and 320 mM KCl (colored lines), compared to reference conditions (black lines; 10 mM Tris and 5% glycerol only). Increasing salt concentration results in shorter diffusion times, indicating more compact AGARP conformations.

Each panel of **Figure 1** shows a comparison between the autocorrelation curves for AGARP in the presence of a high salt concentration (colored) and the reference curves recorded without salt (black). The salt concentrations are chosen so that the ionic strength remains comparable across the panels (∼150 mM for divalent salts, orange; ∼450 mM for monovalent salts, blue). In all cases, the autocorrelation curves shift toward noticeably shorter lag times, which indicates faster diffusion and thus a smaller *R_h_* value of the polyelectrolyte in the presence of salts. However, the degree of this salt-induced protein compaction is significantly greater for the divalent salts, as evidenced by the larger shifts to the left on the logarithmic lag time axis.

As a robustness check, we performed analogous measurements using semi-analytical SEC as a complementary method. SEC was conducted at 10 °C with detection at 215 nm. Each protein sample, containing AGARP at 4 µM, was incubated for 30 minutes in TG buffer containing appropriate salt concentration. Prior to each measurement, the column was equilibrated with the corresponding buffer, and the *R_h_* values were determined with experimental uncertainty from the calibration curve (**Figure S4**).^31^

Consistent with the FCS observations, higher salt concentrations led to an increase in the elution volume (*V_e_*) of the protein in the SEC chromatograms (**Figure 2**), indicating a more compact conformation and thus longer retention within the porous matrix of the column. SEC, in particular, enabled the measurement of the *R_h_* values at higher salt concentrations - up to 2 M NaCl - than was possible using FCS, due to changes in the optical properties of the salt solutions. This revealed saturation of the electrostatic repulsion attenuation effect. At these elevated salt levels, the protein sizes remained largely unchanged regardless of the ion valency.

**Figure 2.**
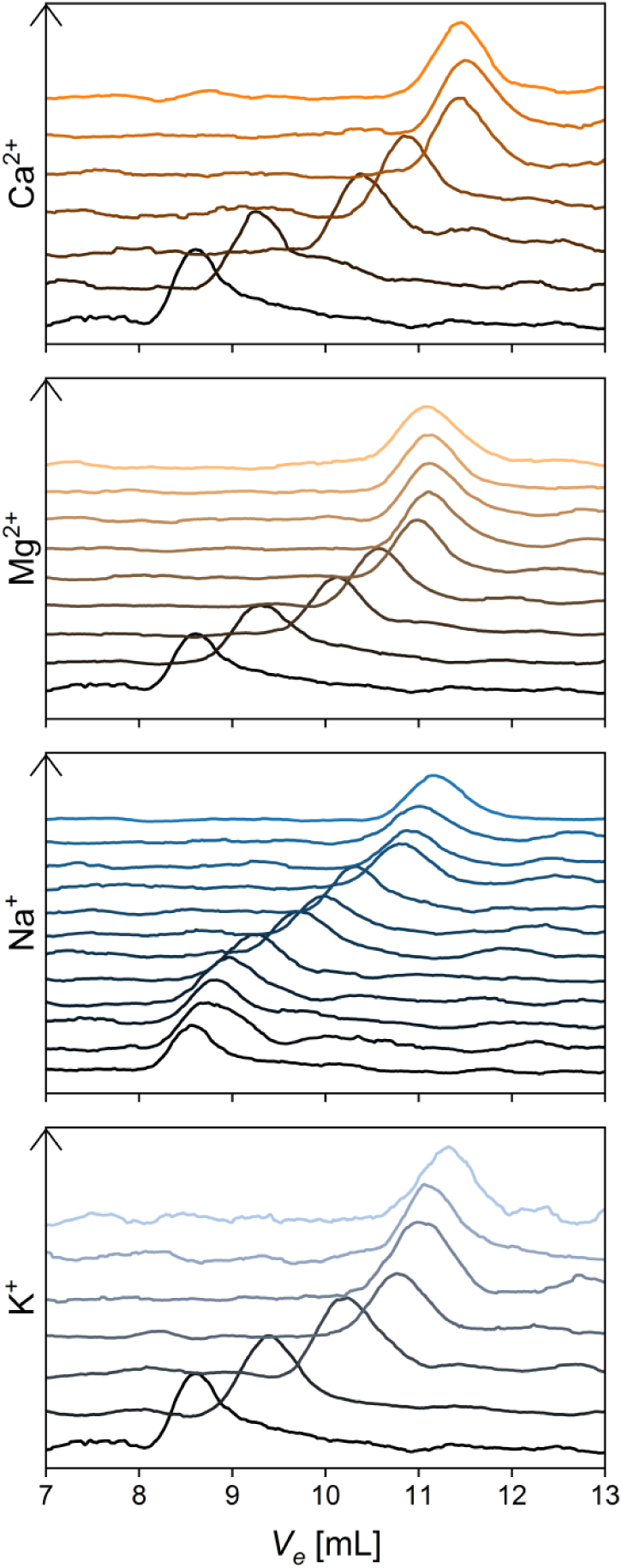
Effects of cations on the diffusion of AGARP, measured by SEC. Absorbance signals recorded at 215 nm for runs at increasing salt concentrations (CaCl_2_, MgCl_2_, NaCl, and KCl) shown staggered for clarity, with the lowest concentration at the bottom (black) and the highest at the top (colored accordingly with **Figure 1**.). Increasing salt concentration results in larger elution volumes, *V_e_*, indicating slower migration through the column matrix, and thus more compact AGARP conformations.

However, the most salient feature reflecting the strength of interactions between the polyanionic protein chain and counterions is the characteristic range of ion activity at which the compaction occurs. Strikingly, the decrease in the hydrodynamic dimensions observed by FCS was triggered by the divalent cations at the activity ranges that were, on average, approximately 25-fold lower than those for the monovalent cations (∼10-fold lower as seen by SEC). These effects point to more specific interactions between the negatively charged protein chain and Ca^2+^ or Mg^2+^, since it cannot be explained simply by the higher ionic strength, which, for divalent ions, is multiplied by three.

The experimental results were compared against two types of *in silico* numerical approaches: CG MD simulations and direct Monte Carlo (MC) sampling (**Figure 3**). AGARP conformations can range from compact to highly extended, which would require an exceptionally large solvation box, thereby rendering all-atom simulations impractical. To address this limitation, we employed the CG model proposed by Dignon *et al.*,^38^ in which protein conformations are parameterized by the positions of Cα atoms that interact *via* two types of potentials: screened electrostatic and modified Lennard-Jones. The electrostatic interactions depend on the product of the charges of the interacting residues (Arg, Lys: +1; Asp, Glu: −1; His: +0.5; and 0 for other residues), their separation and the Debye screening length, which is computed from the ionic strength of the solution. The modified Lennard-Jones potential corresponds to van der Waals forces and hydrophobicity, where the interaction strength and interaction range are computed for each pair of residues^38^ with further temperature-dependent refinements^39^ (see Supplementary Materials for details). Both types of interactions are truncated at a distance threshold.^38^ Simulations were conducted using the user-defined potentials feature of the GROMACS package^41^ for at least 150 microseconds to ensure full equilibration and to obtain precise estimates of both the *R_g_*and *R_h_* values (**Figure 3A**, solid lines). Such a long simulation period is required due to the substantial thermal fluctuations in the instantaneous *R_g_*, which can vary by as much as 50% (**Figure 3A**, translucent lines).

**Figure 3.**
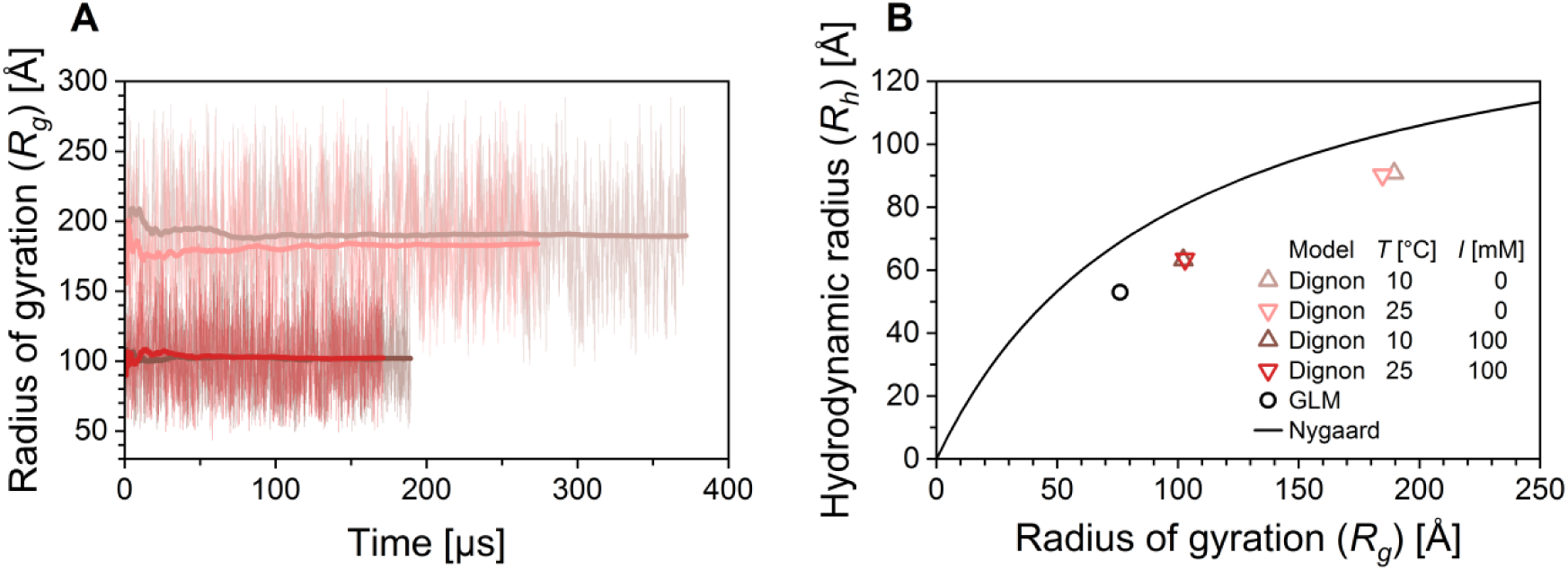
Results of CG MD simulations of AGARP at low (0.001 mM, denoted to as “0”, pastel colors) and high (100 mM, saturated colors) ionic strength conditions, at 10 °C (brown) and 25 °C (red). (A) The instantaneous *R_g_* is plotted against simulation time, along with the cumulative average (thicker line). Despite large fluctuations in *R_g_*, a stable long-time average is achieved. (B) Ensembles of conformers from CG MD (red and brown triangles) and MC GLM (black circle) methods were post-processed using MDA,^36,37^ to calculate *R_h_*; the resulting *R_h_*–*R_g_*pairs were compared with the relationship proposed by Nygaard *et al*.^42^ (black curve). The deviation is attributed to more extended configurations of a larger and more highly charged protein than those included in the training data set.^42^

The second method of conformer generation was the GLM, as described previously,^36^ which operates under the assumption of perfect screening. In this approach, AGARP was modeled as a completely unstructured “linker” chain using a self-avoiding random walk with a steric exclusion range corresponding to the Cα distance.

The ensembles of conformers obtained from both methods were further processed to obtain *R_g_*, and then to calculate *R_h_* using the MDA approach^37^ with the hydrodynamic parameters of chain monomers as described earlier^36^ (**Figure 3B**). The *R_h_ vs*. *R_g_* pairs obtained were then compared with the predictions proposed by Nygaard *et al.*^42^ The origin of the deviations of our results from their *R_h_ vs*. *R_g_* relationship is threefold. First, AGARP is an extremely charged IDP, surpassing the charge density range of their original training dataset. Second, AGARP is longer (506 residues) than the longest protein in the training dataset (450 residues). Third, in the work of Nygaard *et al*.,^42^ the *R_h_* values were computed in a rigid-body approximation,^43,44^ which is not well-suited for highly flexible proteins. In contrast, our work uses MDA, which is specifically designed for IDPs and has been validated on a wide range of IDPs.^36^ Nevertheless, the Nygaard *et al*.^42^ equation still reproduces the correct trend. The quantitative deviation between the predicted and calculated *R_h_* values can be considered small enough to be make the equation suitable for converting *R_g_* predictions from polymer models to *R_h_*, to facilitate further comparison of experimental results with simulations using analytical expressions.

The measured *R_h_* values were analyzed using two frameworks: apparent direct binding, and a polymer-theoretic approach (**Figure 4A-C**). In both the FCS and SEC measurements, the *R_h_* values decreased according to a sigmoidal function of salt activity, *a* (**Figure 4A,B**), with plateaus at the extremes of low and high activity. This supports the applicability of a two-state binding model, assuming identical, non-interacting, entropically independent ion-binding sites on a protein molecule and a large excess of ions compared to the number of binding sites. The apparent dissociation constant, *K_d_*, describes the affinity of an ion for a single binding site, while *ρ* is the observable magnitude of the relative hydrodynamic dimension change due to the binding, according to:

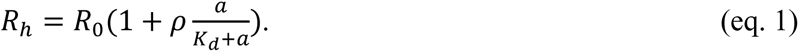

**Figure 4.**
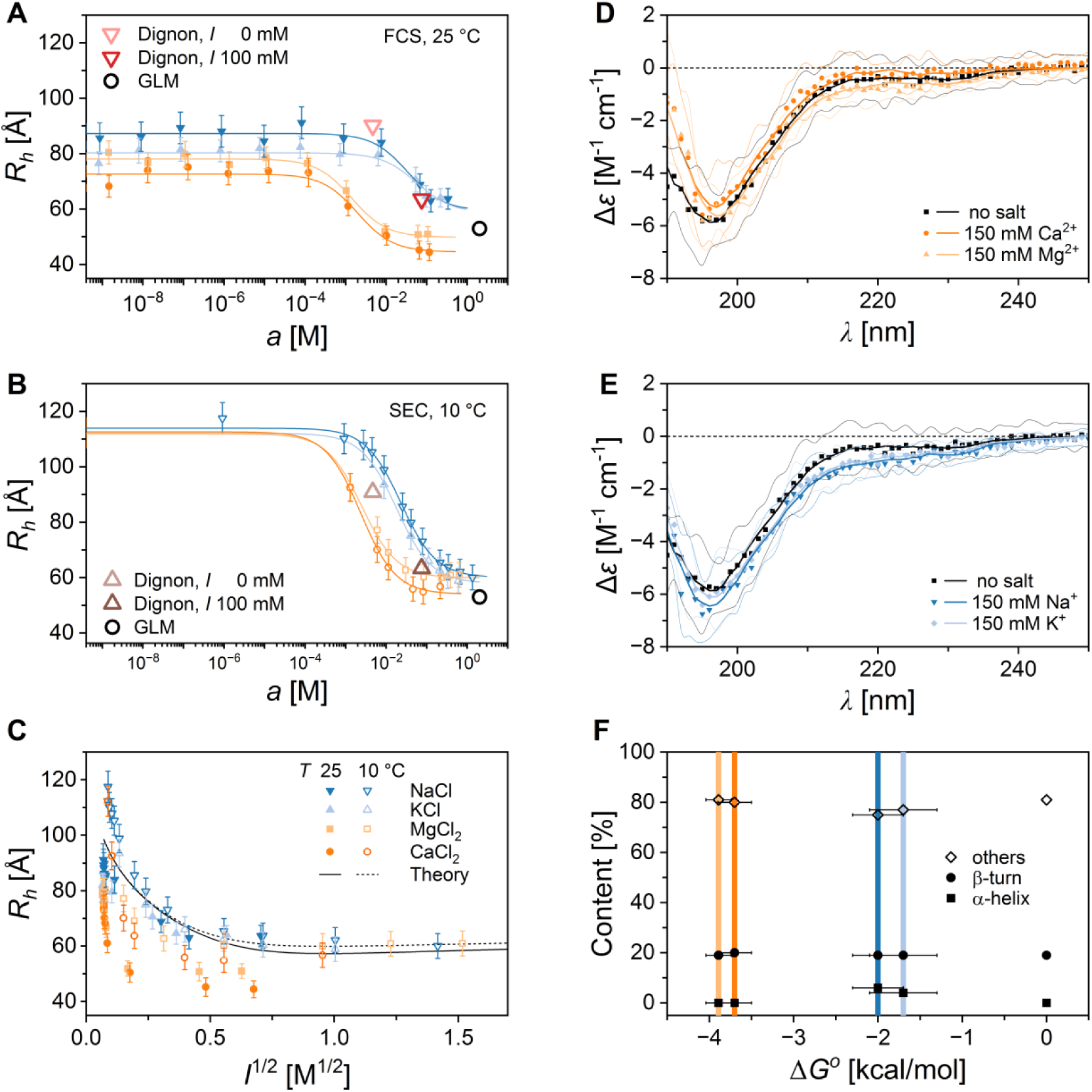
The structure-less electrostatic collapse of AGARP depends on the cation valency. At increasing salt concentrations, the hydrodynamic radius of AGARP, *R_h_*, decreases according to the sigmoidal binding curve (eq. 1, Table 1), as evidenced by (A) FCS and (B) SEC. The *R_h_* derived from CG MD (red and brown triangles) and GLM-MDA (black circle) simulations are in line with the experimental trends for the monovalent salts (blue), whereas the divalent salts (orange) induce AGARP compaction at markedly lower salt activities. (C) Combined FCS and SEC data align well with polymer-based Debye–Hückel screening predictions (Theory, continuous and dotted black curves) for monovalent salts, while significant deviations are observed for divalent cations due to the enhanced compaction. Normalized CD spectra of AGARP in the presence of (D) divalent and (E) monovalent salts, analyzed using BeStSel^47^. (F) Results of CD analysis showing negligible α-helix content regardless of the cation type and Gibbs free energy of the binding, Δ*G^O^*(Table 1). The β-type structures are indiscernible from random coil (others) in IDPs.^48^

The decrease in *R_h_*, which is indicative of increased AGARP compactness, occurs at much lower activities of divalent ions compared to monovalent ions. As shown in **Table 1**, the *K_d_* values obtained from FCS and SEC measurements are similar, with differences of one or two standard errors resulting from numerical fitting. The apparent dissociation constants for divalent ions are about several dozen times lower than those for monovalent ions. To facilitate the comparison, the values of *K_d_* were converted into the changes in Gibbs free energy (Δ*G°*) that accompany the ion binding. Strikingly, the Δ*G°* for the Ca^2+^ and Mg^2+^ cations corresponds to that of salt bridge formation in an aqueous environment (3-5 kJ/mol).^45,46^ For the monovalent Na^+^ and K^+^ cations, however, the Δ*G°* is only about three times the thermal energy (*RT*). This suggests that the divalent ions are more tightly bound, while monovalent ions can only be loosely or transiently coordinated.

**Table 1.**
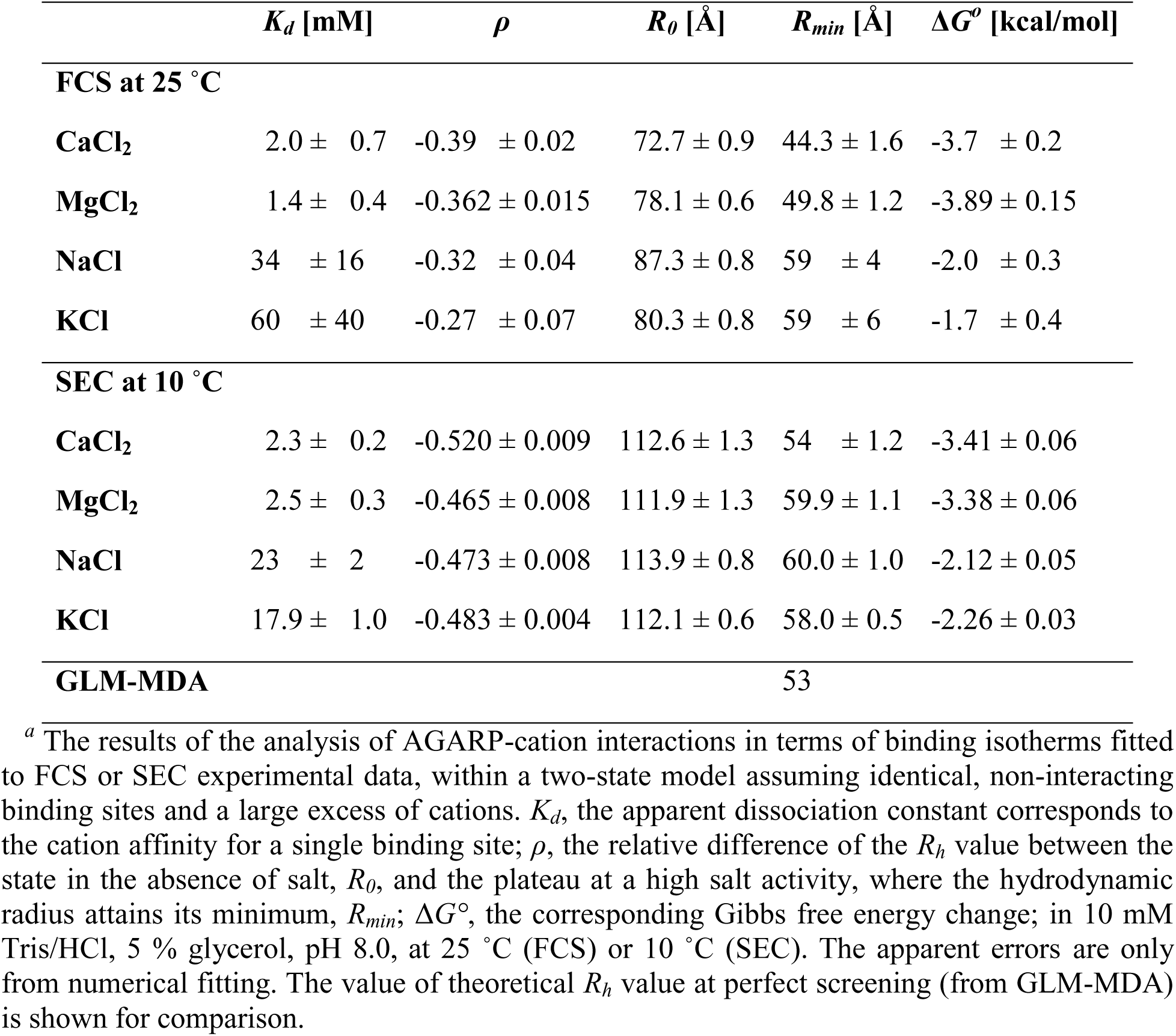
Parameters of Cation Binding by AGARP *^a^*.

While relative changes in the *R_h_* of AGARP upon interaction with ions show little difference between different ions of the same valency using a given method, some differences that reflect the polyelectrolytic nature of the protein are observed between the methods. The relative change measured by SEC appears greater than that observed by FCS. This is primarily driven by the differences at very low salt concentrations, where AGARP adopts highly extended conformations that can affect the reliability of measurements using the Superdex 200 Increase 10/300 GL SEC column, which contains a porous matrix optimized for globular proteins. However, as the chains adopt more compact shapes, this discrepancy disappears. The lower *R_h_* limits, *R_min_*, derived from both methods are in excellent agreement and align closely with the GLM-MDA prediction (**Table 1**, **Figure 4A,B**).

Next, we compare the experimental results with the model (**Figure 4C**) applied by Müller-Späth *et al*.^21^ based on the polymer theory of Higgs and Joanny^40^ that uses Gaussian displacement statistics to describe the screened electrostatic interaction energy of a chain containing *N* monomers and a Kuhn length, *b*. The residues are positively or negatively charged with probabilities *f* or *g*. This interaction can be either repulsive or attractive, and its strength depends on the net absolute charge per residue, *f* + *g*, and the net charge per residue, *f* – *g*. The second type of interaction considered is the effective excluded volume interaction controlled by the parameter *ν*, which is the only fitting parameter of the model. The screening length, *λ_D_*, is assumed to follow the Debye-Hückel dependence on ionic strength, *I*, and the Bjerrum length, *l_B_*, which combines the properties of the liquid and temperature.

The polyampholyte model is described by the following relations:

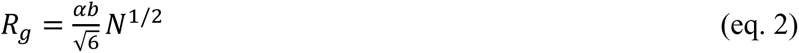

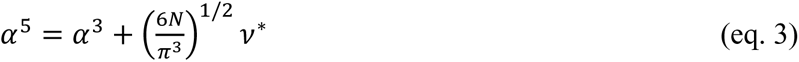

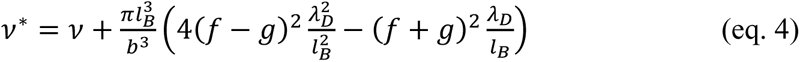

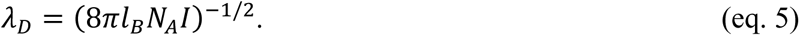

In the presented cases, assuming 2 < *α* < 5 and omitting *α*^3^ term in eq. 3 leads to an approximation error of at most 5% when determining *R_g_*. The values of the constants are listed in the Supplementary Material (**Table S1**). The *R_g_* predictions were combined with the phenomenological relationship established by Nygaard *et al.*^42^ to determine *R_h_*:

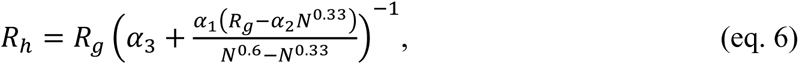

with *α_i_* parameter values as described therein. The combined eqs. 2–5 and 6 were used in the fitting procedure to determine *ν* from either FCS and SEC measurements with monovalent salts, which resulted in similar values of *ν* = 2.42 and *ν* = 2.87, respectively.

**Figure 4C** shows that the polyampholyte theory model adequately describes the electrostatic compaction of AGARP induced by screening by monovalent ions. However, it fails to account for the stronger collapse upon interactions with divalent ions. This further supports the possibility of tight binding in the latter case.

To test whether the observed ∼40% decrease in the hydrodynamic dimensions of AGARP (**Table 1**) is linked to the protein folding, CD spectra were recorded in a TG buffer supplemented with 150 mM salts and without added salt. The CD spectra of AGARP in the presence of di-(**Figure 4D**) and monovalent (**Figure 4E**) cations are almost identical to those under salt-free conditions, displaying only slight changes at 190 and 218 nm, respectively, that are hardly interpretable in terms of the quantitative contributions of secondary structure elements. The BeStSel^47^ analysis revealed that cations, regardless of valency, do not induce the formation of measurable amounts of α-helices (**Figure 4F, Table S2**), the only type of secondary structure that can be reliably quantified by CD in the case of IDPs.^48^ We thus conclude that AGARP compaction caused by both Debye-Hückel screening of electrostatic repulsion and additional enhancement by chelation of divalent ions is a conformational collapse that preserves the disordered state of the protein.

Our results are in line with the previously reported changes in IDP sizes due to screening-induced collapse for the polyelectrolytic IDP prothymosin α^21^ in the presence of monovalent salts. Similarly to AGARP, the collapse of prothymosin α can be described by polymeric theories of Debye screening. The amplitude of this effect (30%), however, is smaller than that for AGARP (40%), despite the slightly higher charge density of prothymosin α. *In silico* studies have also shown that prothymosin α responds selectively to divalent ions,^27^ but this result has yet to be confirmed experimentally. Collapse under the influence of divalent ions, without accompanying secondary structure formation, analogous to AGARP, has been demonstrated experimentally for other polyanionic proteins associated with biomineralization, such as the otolith matrix macromolecule-64 (OMM-64)^30^ and Starmaker.^29^ Although these proteins are similar to AGARP in terms of chain length (608 and 593 residues for OMM-64 and Starmaker, respectively) their smaller charge density (−0.27 and −0.23 e/res, respectively) results in a decreased electrostatic collapse amplitude (∼12% and 30%, respectively) compared to AGARP. These characteristics of polyelectrolytes contrast with the features of polyampholytic proteins, such as, *e.g.*, the basic helix–loop–helix family,^23^ exhibiting the opposite effect of screening-induced swelling at relatively low, physiological monovalent salt concentrations.

Guided by the observations of the structure-free, counterion-induced collapse of AGARP, we propose the following schematic representation of this phenomenon for acid-rich proteins (**Figure 5**). The absence of salt favors extended protein conformations with high *R_h_*values. As the salt concentration increases, a cloud of counterions forms around the protein, suppressing intra-protein electrostatic repulsion, and resulting in relaxed, more compact conformations described by lower *R_h_*values. The strength of this effect is captured by the Debye screening length for monovalent salts. On the other hand, the screening effect, which is three times stronger for divalent salts at the same molar concentration, is insufficient to explain the dramatic collapse of *R_h_*triggered by divalent salts. We propose that specific binding is necessary to explain the favoring of more compact conformations in the presence of Ca^2+^ or Mg^2+^. Moreover, we show that, for natively unstructured acid-rich proteins, this compaction is a process of structure-less collapse. This is supported by two observations: first, CD spectra show a negligible amount of the α-helix content (**Figure 4D-F, Table S2**); second, the minima of the *R_h_* values under both divalent and monovalent salt conditions align with the GLM-MDA’s self-avoiding random walk predictions (**Table 1**).

**Figure 5.**
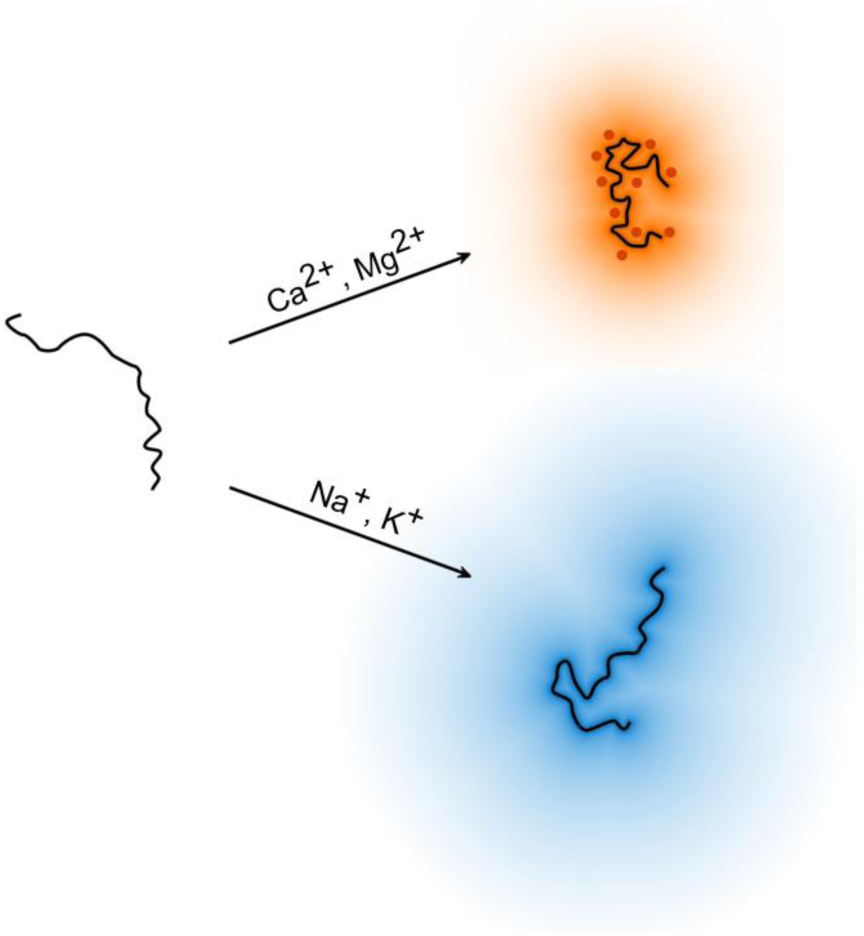
Schematic representation of the enhancement of the electrostatic collapse of an acid-rich protein induced by divalent cations. In a solution of low ionic strength (left), AGARP (black line) adopts extended conformations due to intra-protein repulsion. In the presence of monovalent cations, the repulsion is screened by the surrounding cation cloud (bottom right; blue gradient). In contrast, divalent cations bind more specifically, yielding more effective charge neutralization (top right; red points, orange gradient).

In this work, using the large polyanionic IDP AGARP as a model, we provide both experimental and theoretical evidence that unstructured proteins can interact specifically with counterions, distinguishing between their valencies. We demonstrate additional chelating protein behavior in the presence of divalent cations such as Ca^2+^ and Mg^2+^, in contrast to the simple electrostatic screening observed for monovalent cations like Na^+^ and K^+^. Even upon saturation with counterions, AGARP does not develop secondary structures. We conclude that the IDP undergoes a structure-less electrostatic collapse, a conformational compaction driven by suppression of electrostatic self-repulsion without folding. Our findings suggest that a comprehensive understanding of charged IDP–counterion interactions requires moving beyond the traditional conceptual framework of ion clouds, particularly given critical role of divalent Ca^2+^ and Mg^2+^ cations in the biological function of acid-rich proteins in regulating biomineralization.

## ASSOCIATED CONTENT

The **Supporting Information** is available free of charge.

Materials and Methods, Additional Tables (list of parameters, BeStSel CD analysis results, limiting molar conductivity from SEC), and Figures (bioinformatics analysis of AGARP, count rates from FCS, SEC calibration curve, conductivity from SEC) (PDF)

## AUTHOR INFORMATION

## Author Contributions

B.P.K. and A.N. designed and performed the experiments; M.W designed and performed the *in silico* simulations; R.W. designed and performed the theoretical calculations; B.P.K., R.W., M.W., and A.N. analyzed, interpreted, and visualized the data; B.P.K., R.W., and A.N. wrote the manuscript, with contributions from the other authors; A.M. did preliminary experiments; A.N. acquired the funding and conceived, designed, and supervised the project. All authors have approved the final version of the manuscript.

## Notes

The authors declare no competing financial interest.

## ACKNOWLEDGMENT

We thank Prof. Joanna Trylska for the access to the CD laboratory at the Centre of New Technologies, University of Warsaw. The work was supported by Polish National Science Centre grant no. 2016/22/E/NZ1/00656. The research was performed in the NanoFun laboratories co-financed by ERDF within the Innovation Economy Operational Program POIG.02.02.00-00-025/09.

