## Supplementary Material for "Electrostatic Collapse of Intrinsically Disordered Acid-Rich Protein Is Sensitive to Counterion Valency"

### Materials and Methods

#### *Materials*

The following reagents of analytical grade were used: HCl, KCl (Chempur); ethanol (Pol-Aura); $CaCl_2$, glycerin, $MgCl_2$, NaOH, Triton X-100 (Carl Roth); NaCl (Stanlab); $NaH_2PO_4$, Bovine Serum Albumin (BSA) with monomer fraction ≥ 97% (Sigma-Aldrich); Tris(hydroxymethyl)aminomethane (Tris) (Millipore).

#### *Bioinformatics*

The secondary structure and spatial conformation of AGARP were predicted using AlphaFold2[1] *via* the ColabFold platform.[2] Per-residue confidence scores were computed using the predicted local-distance difference test (pLDDT), based on the method described in Ref.[3]. The schematic representation of the top-ranked model of AGARP (corresponding to the highest pLDDT, **Figure S1**) was drawn using Discovery Studio v19.1.0.18287 (Dassault Systèmes, San Diego). Disorder propensity was predicted with IUPred3.[4]

#### *Buffers*

Water for experiments was purified using a Hydrolab deionizer HLP 5 and Biopak® Polisher. All buffers and demineralized water (except salt stock solutions) were treated with Chelex® 100 resin (Bio-Rad) to remove divalent ions. Stock solutions of NaCl, KCl, $MgCl_2$ and $CaCl_2$ were prepared in water for FCS experiments and in TG buffer (10 mM Tris/HCl, 5% glycerol pH 8.0) for circular dichroism (CD) measurements (25 °C) and size exclusion chromatography (SEC) (10 °C). Salt stock solutions concentrations for FCS were prepared assuming the droplet evaporation rate of 7 µL/11 min. Blocking buffer (20 mM Tris/HCl, 150 mM NaCl, 0.1% (w/v) Triton X-100, 3% (w/v) BSA, pH 8.0) was used for the glass slide passivation procedure.

#### *Protein sample preparation for FCS*

Purified AGARP sample[5] was incubated with Chelex® 100 resin for two hours. Deionized AGARP was labeled with AF488 NHS ester (Lumiprobe) according to the manufacturer's instructions. In short, the protein was incubated in 100 mM phosphate buffer, pH 8.5 with an 8-fold molar excess of dye overnight at 10 °C with gentle stirring. Unbound AF488 was removed by passing the sample twice through a SEC column (Zeba™ Spin Desalting Column 7K MWCO, Thermo Fisher Sci.). Prior to chromatography, the column was rinsed with deionized water until there were no compounds absorbing in the 230-570 nm range in the filtrate. AGARP sample was subsequently dialyzed against TG buffer. A protein concentration of approx. 100 nM was used for FCS experiments.

*FCS measurements*

FCS measurements were performed on an Axio Observer LSM 780 Zeiss inverted confocal microscope (40x/1.2 water immersion objective) with a ConfoCor3 kit. MBS 488 beam splitter and a BP 495-555 emission filter was used. An argon laser was used as the source of the excitation beam. Emitted photons were detected using an avalanche photodiode. The surface of glass slide was passivated with BSA before use. Standard procedure of passivation was applied: the glass slide was sonicated in an ultrasonic cleaner for 2 hours, rinsed with 96% ethanol, blow dried with nitrogen, incubated for 2 hours with 100 μL of evenly distributed blocking buffer, rinsed with water and dried overnight in a desiccator.

Before each experiment the argon laser power, pinhole and objective ring were calibrated with freely diffusing 10 nM AF488 dye solution in deionized water.

FCS titration experiment was performed at 25°C: (1) 30 μL of protein sample in TG, filtered through a 0.22 μm pore size membrane, was incubated for 6 minutes, (2) fluorescence intensity fluctuations were measured 50 times, for 6 seconds each, and (3) 7 μL of salt stock solution in water was carefully added to the AGARP sample. Steps (1)-(3) were repeated for the same sample until the desired final salt concentration was reached (490 mM, 320 mM, 150 mM and 130 mM for NaCl, KCl, $MgCl_2$ and $CaCl_2$, respectively). The same procedure was applied for 10 nM AF488 solution in TG in order to determine the diffusion time of unbound AF488.

*FCS analysis*

The analysis was performed using Zen2010 software (Zeiss), essentially as described previously.[6,7] Count rate traces that reached equilibrium and showed no spikes were selected for further analysis (**Figure S2, S3**). Models of 3D diffusion including a triplet state and either one- or two-components were globally fitted to autocorrelation curves obtained for AF488 and AGARP samples, respectively. In either case, the triplet relaxation times of 4 μs for free AF488 and 2.4 μs for AF488 covalently attached to AGARP, which were obtained in an independent experiment conducted at higher laser power, were fixed. The structural parameter was determined during calibration on AF488 in water and was subsequently fixed for all other curves measured on the same glass slide. For models fitted to protein sample curves, the second component's diffusion time was fixed to that of AF488 measured in TG buffer at the corresponding salt concentration.

The hydrodynamic radius $R_h$ of AGARP was determined from:

$$R_h = \frac{k_B \cdot T \cdot \tau_{AGARP,buf}}{6\pi \cdot D_{AF488,H2O} \cdot \eta_{H2O} \cdot \tau_{AF488,buf}} \qquad \text{(eq. S1)}$$

where $k_B$ - Boltzmann constant, $T$ - temperature (25 °C), $\tau_{AGARP,buf}$ and $\tau_{AF488,buf}$ - diffusion time of AGARP and AF488 in TG with corresponding salt concentration, respectively, $D_{AF488,H2O}$ - diffusion coefficient of AF488 at 25 °C, calculated from its value at 22.5 °C,[8] using Stokes-Einstein relation, assuming a constant ratio of temperature to viscosity and $\eta_{H2O}$ - water viscosity at 25 °C taken from https://wiki.anton-paar.com/en/water/.

*SEC*

SEC was conducted using a Superdex 200 Increase 10/300 GL column (Sigma) at 10 °C with a flow rate of 0.5 mL/min. The injection volume was 100 µL, and detection was performed at 215 nm. Prior to each experiment, the protein sample at 4 µM was incubated for 30 minutes in TG buffer containing the appropriate salt concentration, and the column was equilibrated with the same buffer.

To determine the $R_h$ of AGARP, the system was calibrated using standard proteins in TG with 100 mM NaCl: lysozyme (19 Å),[9] α-chymotrypsinogen A (22.4 Å),[10] BSA monomer (35.5 Å)[10] and dimer (43 Å),[10] and apoferritin monomer (60.3 Å)[11] and dimer (88.8 Å).[11] An exponential function described by the equation:

$$R_h = (Y_0 - Plateau) \cdot \exp(-A \cdot K_{AV}) + Plateau \qquad \text{(eq. S2)}$$

where $Y_0$, *Plateau* and $A$ are fitting parameters and $K_{AV}$ is the partition coefficient derived from SEC, was fitted to calibration data points (**Figure S4**).

A bi-Gaussian function was fitted to AGARP SEC curves.

To verify that the salt concentration remained at the desired level during the SEC run, despite the use of a method commonly employed for desalting, conductivity ($\kappa$) measurements were analyzed. As the conductivity remained stable throughout the run (**Figure S5**), an average value was calculated and analyzed as a function of salt concentration ($c$) for each type of salt

(**Figure S6**). To confirm the accuracy of the salt concentrations, Kohlrausch's Law for strong electrolytes was fitted to the data:

$$\kappa = \Lambda_m^0 \cdot c - K \cdot c^{3/2} \quad \text{(eq. S3)}$$

where $\Lambda_m^0$ – limiting molar conductivity, $K$ – empirical constant.

*Calculations of salt activity*

For FCS, the actual concentrations of salts, Tris, and glycerol in the droplet were estimated based on the fitted number of particles, the number of moles of each component added, and the initial droplet volume.

Ionic strength ($I$) was calculated for each solution, including contributions from all ionic species (salt ions and $TrisH^+$). The mean activity coefficient ($\gamma_\pm$) was then obtained from a $\gamma_\pm$ versus $I$ curve, constructed based on data from.[12] Finally, the mean electrolyte activity ($a$) was calculated using the relationship[13]:

$$a = c_\pm \cdot \gamma_\pm \quad \text{(eq. S4)}$$

where $c_\pm$ denotes the mean molar concentration of the electrolyte[13]:

$$c_\pm = c(x^x \cdot y^y)^{1/\nu} \quad \text{(eq. S5)}$$

where $c$ – molar electrolyte concentration, $x$, $y$ – number of moles of cations and anions, respectively, per formula unit of the salt and $\nu$ – total number of ions produced by the dissociation of one formula unit of salt ($\nu = x + y$).

*Two-state ion binding model*

A binding model (eq. S6) assuming identical, non-interacting, and entropically independent binding sites, was fitted to the experimental data points, according to the equation:

$$R_h = R_0 \cdot \left(1 + \rho \frac{a}{K_d + a}\right) \quad \text{(eq. S6)}$$

where $R_0$ is $R_h$ of AGARP in the absence of a given salt (buffer only), $K_d$ – apparent dissociation constant per one ion-binding site, and $\rho$ – relative change of the $R_h$ between the plateaus in the limits of low ($R_0$) and high ($R_{min}$) salt concentrations, similarly as in the works[14–16]:

$$\rho = \frac{R_0 - R_{min}}{R_0} \quad \text{(eq. S7)}$$

The Gibbs free energy change ($\Delta G^o$) was calculated from the standard thermodynamic equation:

$$\Delta G^o = RT \ln K_d \quad \text{(eq. S8)}$$

where $R$ – gas constant, T – absolute temperature.

*CD spectroscopy*

CD spectra were recorded using a MOS-450/AF-CD spectrometer (Bio-Logic). A xenon lamp served as the light source. Far-UV CD spectra were measured in the wavelength range of 190–260 nm using a 0.1 mm quartz cuvette at 25 °C. Samples consisted of 14.1 µM AGARP in pure TG, and 12.5 µM AGARP in TG with the addition of NaCl, KCl, $MgCl_2$, or $CaCl_2$ at a concentration of 150 mM. Measurements were performed immediately after salt addition, with four replicates per sample. The CD signal was collected for 10 seconds at each nanometer. Scans were averaged, and the contribution from the corresponding buffer was subtracted from the AGARP spectra.

*Electrostatic expansion model*

We compare the experimental results with the model presented by Müller-Späth *et al.*[17] based on polymer-theory considerations. The theoretical calculation of the radius of gyration ($R_g$) is based on Higgs and Joanny[18] and uses Gaussian displacement statistics to describe screened electrostatic interaction energy (with screening length $\lambda_D$) of chain with $N$ monomers which are assumed to be positively (negatively) charged with probability $f$ ($g$). This interaction can be either repulsive or attractive and its strength depends on either net charge per residue $f + g$ or net absolute charge per residue $f - g$. Second type of interaction taken into account is the effective excluded volume interaction controlled by the parameter $v$ which is the only fitting parameter of the model. The screening length $\lambda_D$ is assumed to follow Debye-Hückel theory dependence on ionic strength $I$ and Bjerrum length $l_B$ which combines properties of the liquid and temperature.

The model is described by the following relations:

$$R_g = \frac{\alpha b}{\sqrt{6}} N^{1/2} \quad \text{(eq. S9)}$$

$$\alpha^5 = \alpha^3 + \left(\frac{6N}{\pi^3}\right)^{1/2} \nu^* \qquad \text{(eq. S10)}$$

$$\nu^* = \nu + \frac{\pi l_B^3}{b^3}\left(4(f-g)^2 \frac{\lambda_D^2}{l_B^2} - (f+g)^2 \frac{\lambda_D}{l_B}\right) \qquad \text{(eq. S11)}$$

$$\lambda_D = (8\pi l_B N_A I)^{-1/2} \qquad \text{(eq. S12)}$$

In the presented cases $2 < \alpha < 5$ and omitting $\alpha^3$ term in eq. S10 leads to at most 3% approximation error in determining $R_g$.

The $R_g$ predictions were combined with a phenomenological relationship established by Nygaard *et al.*[19] to determine $R_h$:

$$R_h = R_g \left(\alpha_3 + \frac{\alpha_1(R_g - \alpha_2 N^{0.33})}{N^{0.6} - N^{0.33}}\right)^{-1} \qquad \text{(eq. S13)}$$

The parameters of the eq. S13 were determined based on a fitting procedure where the relationship between $R_h$ and $R_g$ was studied *in silico*. Therein conformers of IDPs were generated and $R_h$ was computed assuming rigid body motion in Stokes flow.

The combined equations S9-S12 were used in the fitting procedure to determine the excluded volume parameter $\nu$ from either FCS and SEC measurements in monovalent salts. Both of these fitting procedures resulted in similar values $\nu = 2.42$ and $\nu = 2.87$, respectively.

*Numerical simulations*

We performed numerical simulations to obtain equilibrium ensembles of AGARP conformers using two methods: coarse grained molecular dynamics using the force field proposed by Dignon *et al.*[20] later referred to as Dignon model, and direct sampling, called globule-linker-model (GLM), in perfect screening regime using a method we proposed earlier.[7]

Dignon force field for molecular dynamics attempts to solve several problems. The classical full-atom force fields were parameterized for structured proteins and are not well suited for IDPs. Nevertheless, some modifications and force field parameterizations have been developed specifically for IDPs.

Second problem is the structural flexibility of IDPs — their conformations are highly diverse, ranging from compact to elongated forms. To efficiently simulate this diversity using a full-atom model, the solvation box must be very large to accommodate both compact and extended IDP structures. This may be feasible for small proteins, but becomes impractical for large IDPs. To overcome this, coarse-grained models have been introduced for simulating large systems. One such model was proposed by Dignon.[20]

Most models make similar approximations: proteins are represented as centers located on Cα atoms, interacting through two types of potentials. The first interaction is electrostatics. Each center is assigned a net charge (Arg, Lys: +1; Asp, Glu: -1; His: +0.5) and zero for other residues. The centers with charges $q_i$, $q_j$ interact *via* a screened Coulombic interaction with a screening length $\lambda_D$ given by Debye-Hückel theory[21]:

$$E = \frac{q_i \cdot q_j}{4\pi D r} \cdot \exp\left(-\frac{r}{\lambda_D}\right) \quad \text{(eq. S14)}$$

with $D$ = 80, the dielectric constant of water. The second interaction type is hydrophobicity, modeled using a modified Lennard-Jones potential:

$$E_{LJ} = 4\varepsilon\left[\left(\frac{\sigma_{ij}}{r_{ij}}\right)^{12} - \left(\frac{\sigma_{ij}}{r_{ij}}\right)^{6}\right] \quad \text{(eq. S15)}$$

The total interaction is thus:

$$E_{ij} = \begin{cases} E_{LJ} + \left(1 - \lambda_{ij}\right) \cdot \varepsilon, & \text{if } r \le 2^{1/6} \cdot \sigma_{ij} \\ \lambda_{ij} \cdot E_{LJ}, & \text{if } r > 2^{1/6} \cdot \sigma_{ij} \end{cases} \quad \text{(eq. S16)}$$

Here, $\lambda_{ij}$ is the interaction strength between amino acids $i$ and $j$, based on their hydrophobic properties. Each amino acid type $i$ has a hydropathy index $\lambda_i$, and $\lambda_{ij}$ is calculated as the arithmetic average $\lambda_{ij} = 1/2(\lambda_i + \lambda_j)$. Similarly, $\sigma_{ij}$ is the arithmetic average of effective radii $\sigma_i$ representing the amino acids. The parameter $\varepsilon$ is fitted to balance both electrostatic and hydrophobic interactions for best results.

Dignon later improved this model to incorporate temperature-dependent modifications of hydropathy.[22] The hydropathy value $\lambda_i$ changes with temperature, and amino acids are divided into five groups, each scaling their parameters individually. Additional modifications include parameters for post-translational modifications such as phosphoserine and phosphothreonine.[23] These models are often used to study phase separation of biomolecules. Their parameterization is typically based on calculations of the radius of gyration ($R_g$) of various IDPs. Thus, it is useful to validate the $R_g$ of our molecule (a large IDP).

We used the GROMACS package[24] for which we implemented the Dignon model. As there is no native implementation of this model in GROMACS, we used simulations with userdefined potentials. Each pair of amino acid types has its own potential file, containing both electrostatic and Lennard-Jones-like interactions. The interaction energies are not calculated using analytical formulas, but are instead read from files. While this approach slows down calculations, it allows for flexibility in defining interaction potentials, and enables the full use of GROMACS features and support for model modifications.

The AGARP conformers were generated by the above CG MD simulations at low (0.001 mM) and high (100 mM) ionic strengths, at temperatures of 10 °C and 25 °C. After omitting the first 1 μs of the simulations, the structures were written down every 5 ns.

*Hydrodynamic radius computation*

The ensembles of conformers from both methods were post-processed to obtain the radius of gyration ($R_g$) and the hydrodynamic radius ($R_h$). The Rg values were calculated for each structure and then averaged. The $R_h$ values were calculated using the Minimum Dissipation Approximation (MDA) proposed by Cichocki *et al.*[25] with the hydrodynamic parameters of the monomer beads as described earlier.[7] Since the $R_h$ value could not be calculated using MDA for all structures, 500 random structures were drawn from each set of conditions, and the $R_h$ value was calculated for the ensemble composed of these structures. The random selection process was repeated 50 times, and then the average $R_h$ was calculated from 50 ensembles. The obtained $R_h$ – $R_g$ pairs were compared with the relationship proposed by Nygaard *et al.*[19]

Because of the extreme charge density of the protein AGARP its configurations deviate from random chain statistics further than earlier training data, resulting in unusually large values of $R_g$ at a given value of $R_h$, or, equivalently, phenomenological function predicted $R_h$ around 20% larger than the value computed directly from the conformers using MDA method. This result can be understood in terms of power means of inter-monomer separations – $R_g$ is a degree 2 power mean, while $R_h$ closely follows degree -1 power mean (cf. Kirkwood-Reisman approximation[26]) thus as configurations become more extended due to electrostatic repulsion $R_g$ grows faster than $R_h$.

### Additional Tables

**Table S1.** List of parameters and their values used in the polyampholyte model.

| Symbol | Source | Value | Description |
|---|---|---|---|
| $R_g$ | c | - | Radius of gyration |
| $\alpha$ | c | - | Expansion factor |
| $b$ | k | 3.8 Å | Segment length |
| $N$ | k | 506 | Number of monomers |
| $v^*$ | c | - | Effective excluded volume parameter |
| $v$ | f | 2.42 or 2.87 | Effective volume parameter (data from FCS and SEC, respectively) |
| $l_B$ | k | 7.1 Å | Bjerrum length |
| $f$ | k | 65/506 | Fraction of positive charges per monomer |
| $g$ | k | 212/506 | Fraction of negative charges per monomer |
| $\lambda_D$ | c | | Debye screening length |
| $N_A$ | k | $6.02 \cdot 10^{23}$ mol$^{-1}$ | Avogadro's number |
| $I$ | e | | Ionic strength |
| $R_h$ | c | | Hydrodynamic radius |
| $\alpha_1, \alpha_2, \alpha_3$ | k | 0.216 Å$^{-1}$, 4.06 Å, 0.821 | Fitted constants[19] |

Abbreviations: c – computed from other quantities, k – known, f – fitted, e – varied experimentally

**Table S2.** Secondary structure content of AGARP in the absence of salt and in the presence of 150 mM KCl, NaCl, $CaCl_2$ and $MgCl_2$ estimated by BeStSel.[27]

| | **α-helix** | **β-turn** | **Others** |
|---|---|---|---|
| no salt | 0 | 19 | 81 |
| $CaCl_2$ | 0 | 20 | 80 |
| $MgCl_2$ | 0 | 19 | 81 |
| NaCl | 6 | 19 | 75 |
| KCl | 4 | 19 | 77 |

**Table S3**. The fitted limiting molar conductivity ($\Lambda_m^0$) in eq. S3 compared with its literature value.[28]

| | **$\Lambda_m^0$ [(mS m$^2$)/mol]** | |
|---|---|---|
| **Salt** | **Fit eq. S3** | **Literature[28]** |
| $CaCl_2$ | 25.0 ± 1.7 | 27.2 |
| $MgCl_2$ | 23.4 ± 0.6 | 25.9 |
| NaCl | 12.4 ± 0.2 | 12.6 |
| KCl | 14.1 ± 0.5 | 15.0 |

The experimental values obtained are in a very good agreement with the literature data. The slight discrepancy results from the fact that the condition $\Lambda_m^0 \gg K\sqrt{c}$, as assumed in Kohlrausch's Law for strong electrolytes, is not fully satisfied. The results prove that the actual ionic conditions in the column matrix during the SEC experiments with AGARP matched the nominal salt concentration values.

### Additional Figures

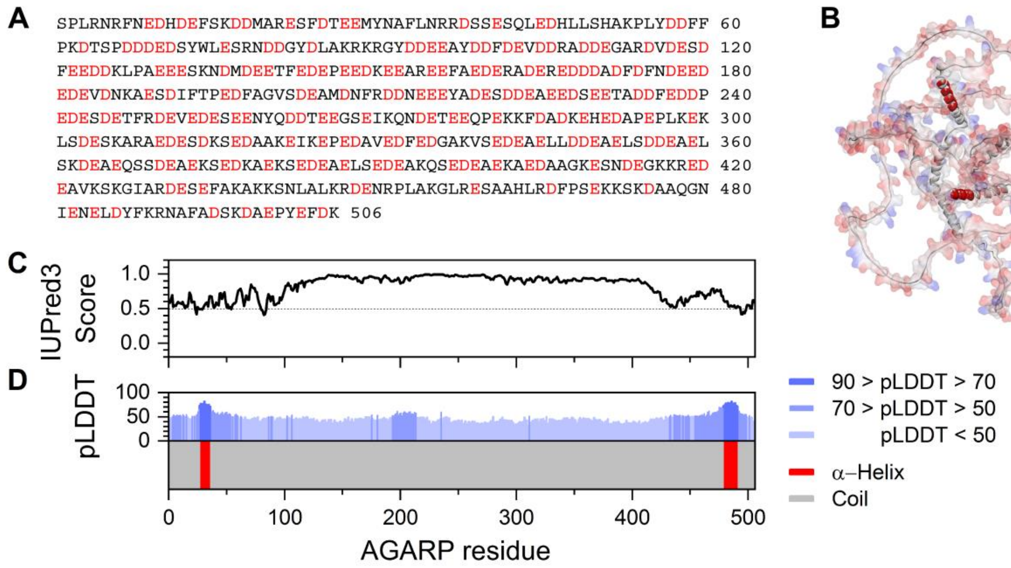


**Figure S1.** Bioinformatics analysis of AGARP. (A) AGARP sequence with acidic amino acid residues marked in red; (B) putative conformation of AGARP chain predicted with AlphaFold2[1,2] with short α-helices shown in red and a coil conformation in grey, presented in a solvent-accessible surface area representation colored by interpolated charge; (C) a disorder propensity obtained using IUPred3[4] with a threshold indicated by a dashed line and (D) a secondary structure predicted by AlphaFold2[1,2] (bottom panel) with the corresponding confidence of prediction (pLDDT) (upper panel). Only the α-helices with pLDDT > 70 are shown in red.

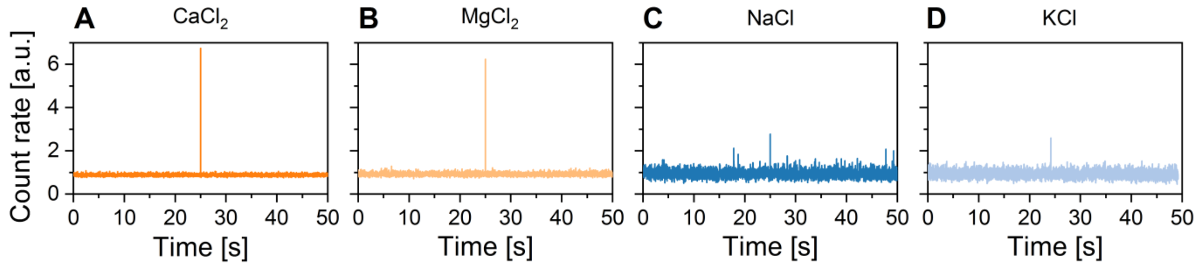


**Figure S2**. Changes in the confocal fluorescence count rate over time, baseline-corrected and normalized to the count rate corresponding to monomeric AF488-labelled AGARP, with the highest measured spikes corresponding to aggregates formed in the presence of (A) 150 mM $CaCl_2$, (B) 8 mM $MgCl_2$, (C) 9 mM NaCl, and (D) 1 nM KCl. The count rate traces containing spikes were discarded from the FCS analysis.

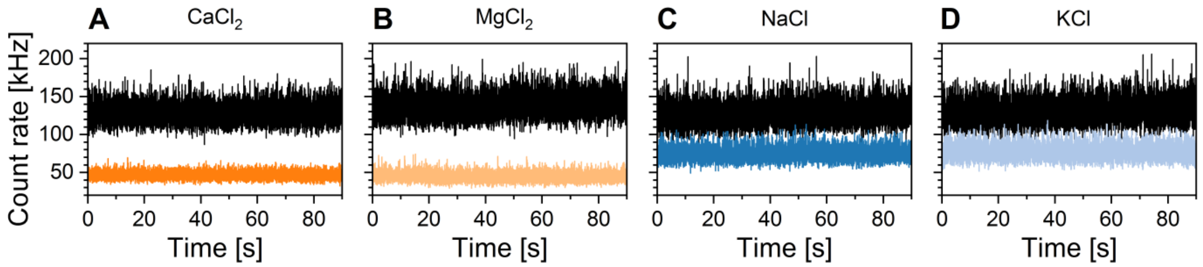


**Figure S3**. Changes in the confocal fluorescence count rate over time for AF488-labelled monomeric AGARP in the absence of salt (black in all graphs) and in the presence of (A) 150 mM $CaCl_2$ (dark orange), (B) 130 mM $MgCl_2$ (light orange), (C) 490 mM NaCl (dark blue) and (D) 320 mM KCl (light blue). The count rate of samples containing salts decreases with respect to salt-free conditions as the addition of constant aliquots of salt solution during titration exceeds the rate of water evaporation. The evaporation was slower at higher salt concentrations.

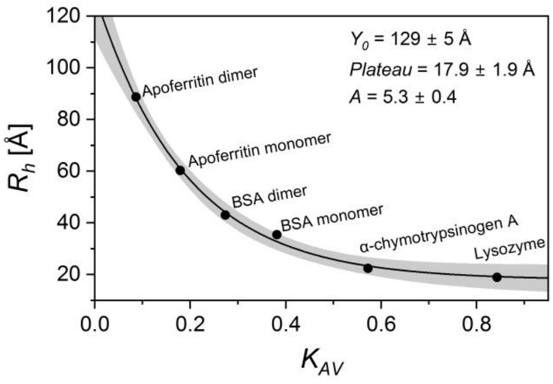


**Figure S4.** SEC calibration curve. Solid line, fitted eq. S2; shaded area, 95% confidence interval.

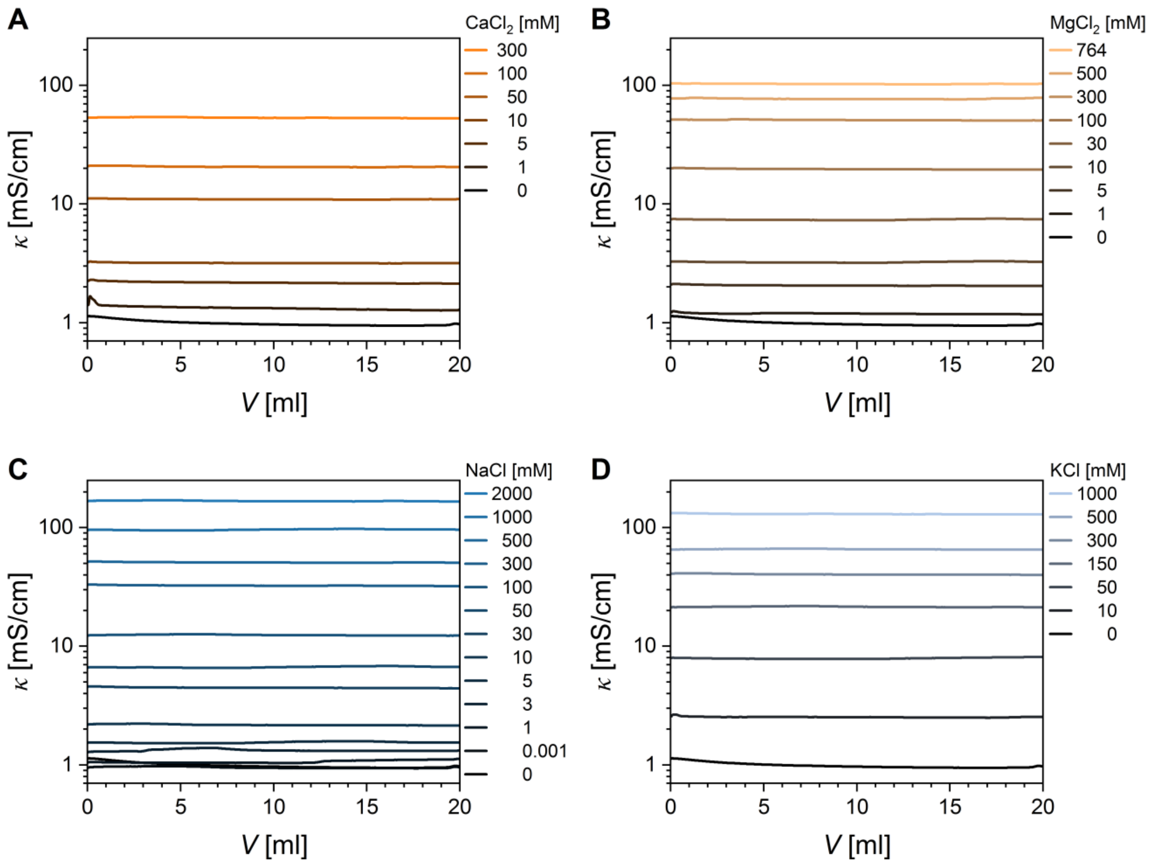


**Figure S5**. Dependence of conductivity (*κ*) on elution volume (*V*) for different concentration ranges of (A) $CaCl_2$, (B) $MgCl_2$, (C) NaCl, (D) KCl during the size exclusion chromatography experiments.

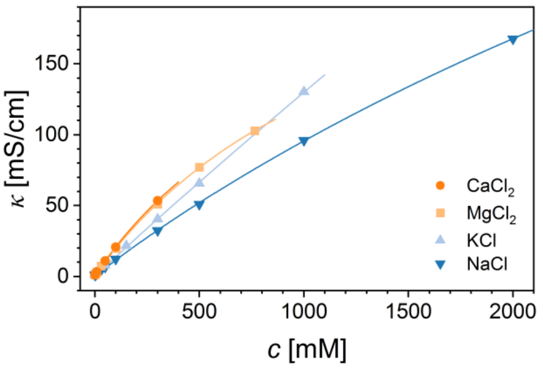


**Figure S6**. Analysis of the dependence of conductivity (*κ*) on salt concentration (*c*). Solid line, fitted eq. S3. Fitted parameters are gathered in **Table S3**.
